# Dexamethasone, a direct modulator of AQP2 in Menière’s disease

**DOI:** 10.1101/2022.01.25.477763

**Authors:** Robin Mom, Julien Robert-Paganin, Thierry Mom, Christian Chabbert, Stéphane Réty, Daniel Auguin

## Abstract

Menière’s disease is a chronic illness characterized by intermittent episodes of vertigo associated with fluctuating sensorineural hearing loss, tinnitus and aural pressure. This pathology strongly correlates with a dilatation of the fluid compartment of the endolymph, so-called hydrops. Dexamethasone is one of the therapeutic approaches recommended when conventional antivertigo treatments have failed. Several mechanisms of actions have been hypothesized for the mode of action of dexamethasone such as anti-inflammatory effect or as a regulator of the inner ear water homeostasis. However, none of them have been experimentally confirmed so far. Aquaporins (AQPs) are transmembrane water channels and are hence central in the regulation of trans-cellular water fluxes. In the present study we investigated the hypothesis that dexamethasone could impact water fluxes in the inner ear through direct interaction with AQP2. We addressed this question through molecular dynamics simulations approaches and managed to demonstrate a direct interaction between AQP2 and dexamethasone and its significant impact on the channel water permeability. We also describe the molecular mechanisms involved in dexamethasone binding and in its regulatory action upon AQP2 function.

**Highlights:**

- AQP2 water permeability is modulated by dexamethasone at physiological concentrations
- The interaction impacts water fluxes through a direct interaction with the extra-cellular surface of the aquaporin
- Key interactions implicate conserved residues of the ar/R constriction
- New insights on corticosteroids mode of actions in Menière’s disease treatment

## Introduction

Menière’s disease (MD), first described by Prosper Menière^1^ is a chronic illness characterized by intermittent episodes of vertigo associated with fluctuating sensorineural hearing loss, tinnitus, and aural pressure^2^.

The main histopathological trait correlating with most MD cases is endolymphatic hydrops (EH), a dilatation of the fluid compartment of the endolymph. It was indeed described by Prosper Menière in 1861, then reported on cadaveric specimen of inner ears from deceased MD patients^3^ and currently visualized in numerous MD patients by means of magnetic resonance imaging of the diseased ear (Hydrops found in 84% of MD ears^4^). This phenomenon is now considered to be a marker of pathology^5,6^, although its direct involvement in the symptoms of the disease remains to be proven^7,8^. The inducing factors of the EH are still unknown. However, it is believed to result from a massive influx of water from perilymph, the liquid compartment adjacent to endolymph^9,10^. Different causes have been put forward to explain this phenomenon, without any being particularly prioritized.

Aquaporins are transmembrane proteins which allow the transfer of water molecules across the biological membranes of most living cells^11–13^. Among them, aquaporin 2 (AQP2) has particularly aroused the interest of researchers due to its role in the control of transmembrane water fluxes^14–16^ and because it is expressed in areas of fluid exchange between perilymph and endolymph^10,15,17–20^. A conformational state of the AQP2 tetramer corresponding to open and functional water channels indeed allows, under the effect of the electrochemical gradient, a transfer of water between the two adjacent fluid compartments of the inner ear. This peculiarity makes it a prime target to explain the phenomenon of EH, in the event of a mutation that would affect the structure or the function of AQP2^21,22^.

Different pharmacological approaches have been developed in an attempt to absorb the accumulation of water responsible for EH. The administration of diuretics is part of the MD patient care recommendations^23^ to reduce hydrops and alleviate the associated vertigo syndrome. Trans-tympanic administration of dexamethasone is also one of the therapeutic approaches recommended in MD, when conventional antivertigo treatments have failed^23^. This approach has shown significant benefits in the treatment of MD^24–27^. Several mechanisms of action have been put forward to explain the effect of dexamethasone in the treatment of MD, such as an anti-inflammatory effect as in the context of acute peripheral vestibulopathy, or a direct modulation of fluid fluxes at the base of the EH formation^26,28–30^. However, none of these hypotheses have so far been substantiated theoretically and verified by experiment.

In the present study, we tested the hypothesis of a direct interaction of dexamethasone with AQP2. Through this interaction, AQP2 water transport activity would be impaired, leading to changes in water fluxes in the inner ear. To test this hypothesis, we addressed the likelihood of this interaction and looked at the underlying molecular mechanisms through molecular dynamics simulations.

## Results

### I. AQP2-DEX interaction site

First, to find a putative interaction site for dexamethasone in AQP2, we used the X-ray structure of the mouse agonist glucocorticoids receptor interacting with dexamethasone (PDB code: 3mne^44^) as a reference (figure 2.A). From this structure, we can notice the protein residues of the interaction site complement very well the amphipathic nature of the dexamethasone. Indeed, we find both polar or charged residues such as asparagine, glutamine or arginine and hydrophobic ones such as isoleucine or tryptophane (figure 2.A). Tryptophane though can also act as an hydrogen bond donor and is hence able to accommodate both polar or hydrophobic parts of the dexamethasone. When we compared this interaction site with the extra-cellular vestibules of AQP2, we found the above mentioned features were conserved. On this part of the AQP, are found an arginine, an asparagine, a glutamine and a tryptophane as well (figure 2.B). Plus, the intrinsic amphipatic nature of the conducting pore^45,46^ characterized by the opposition between polar water molecules interaction sites and hydrophobic slides is leading to very similar features. This amphipathic nature of the AQPs conducting pore allows for very stringent and selective conduction properties^39^ but could also accommodate other amphipatic molecules such as dexamethasone. On top of this similar nature with the native interaction site of dexamethasone, the extra-cellular vestibules of AQPs also present a very conserved residue among the family : the arginine of the aromatic/arginine (ar/R) constriction^11^. This ar/R constriction is named after this arginine and is well known to be determinant in the function of the protein such as it is also called the selectivity filter of the pore. This constriction corresponds to the narrowest part of the channel and its role in the selectivity of the channel has been confirmed by both theoretical studies^47,48^ and mutation experiments^49^. Indeed, the composition of the constriction changes both the size and the hydrophobic nature of the pore which are understood as the two criteria on which relies the selectivity of the AQP ^39^. More recently, several studies further confirmed the critical role of the arginine of the ar/R constriction on the transport properties of the AQP by highlighting significant impact of the position of its side chain in the pore on the water permeability^34,35^. Thereafter, because of the similarities between the extra-cellular vestibules of AQP2 and the interaction site of the dexamethasone and because of the presence of very conserved residues known for their role in transport properties of AQPs, we decided to investigate the ability of these vestibules to fix dexamethasone. In order to do so, we manually placed one dexamethasone molecule on each four extra-cellular vestibules of the X-ray structure of human AQP2 (PDB code: 4nef^50^). We mimicked the interactions observed on the X-ray structure of the mouse agonist glucocorticoids receptor : the oxygen of the carbonyl oxygen is positioned close to the arginine, the hydroxyl groups close to the polar asparagine and glutamine while the hydrophobic side of the dexamethasone faces the hydrophobic part of the channel ending at the extra-cellular surface with the tryptophan (figure 2.A and B).

**Figure 1.**
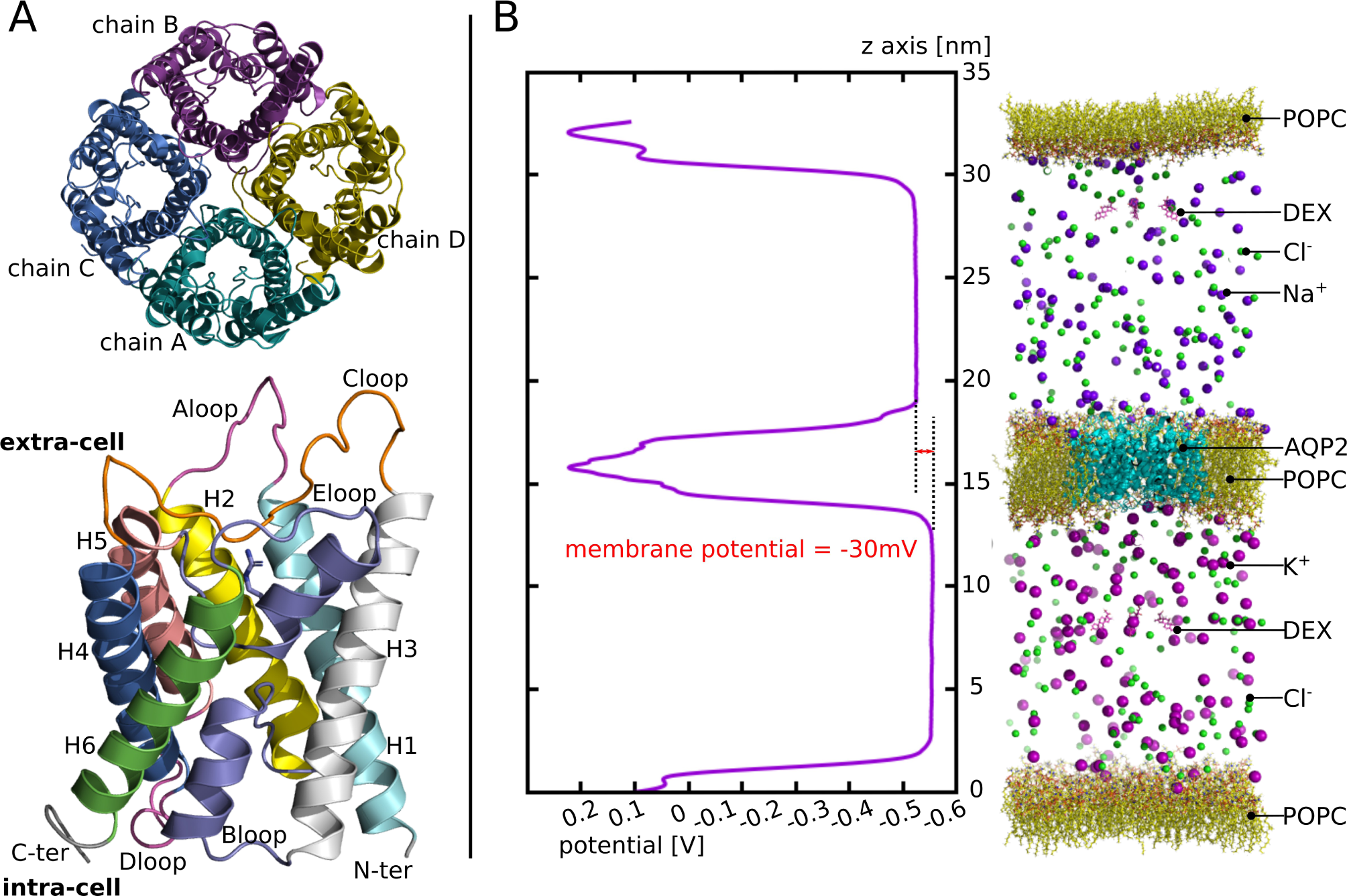
AQP2 structure and illustration of the third experimental setup. **A**. Schematic representation of the tetrameric assembly of AQP2 (on top) and of the structural features of each monomer (on the bottom) extracted from human AQP2 crystallographic structure (PDB code: 4nef). The tetramer is represented as viewed from the extra-cellular compartment and the monomer from a lateral view. On the monomer, Bloop and Eloop colored in purple meet at the center of the monomer. They both incorporate a small alpha helix (HB and HE, not mentioned on the legend). One of them hold the arginine of the ar/R constriction (whose side chain is represented) situated on the extra-cellular half of the channel. **B**. In the left part : the potential along the z axis of the simulation box is displayed. The whole 250 ns of the production phase are taken for membrane potential calculation. In the right part : schematic representation of the atomic system simulated. For clarity purpose water molecules are not represented.

**Figure 2.**
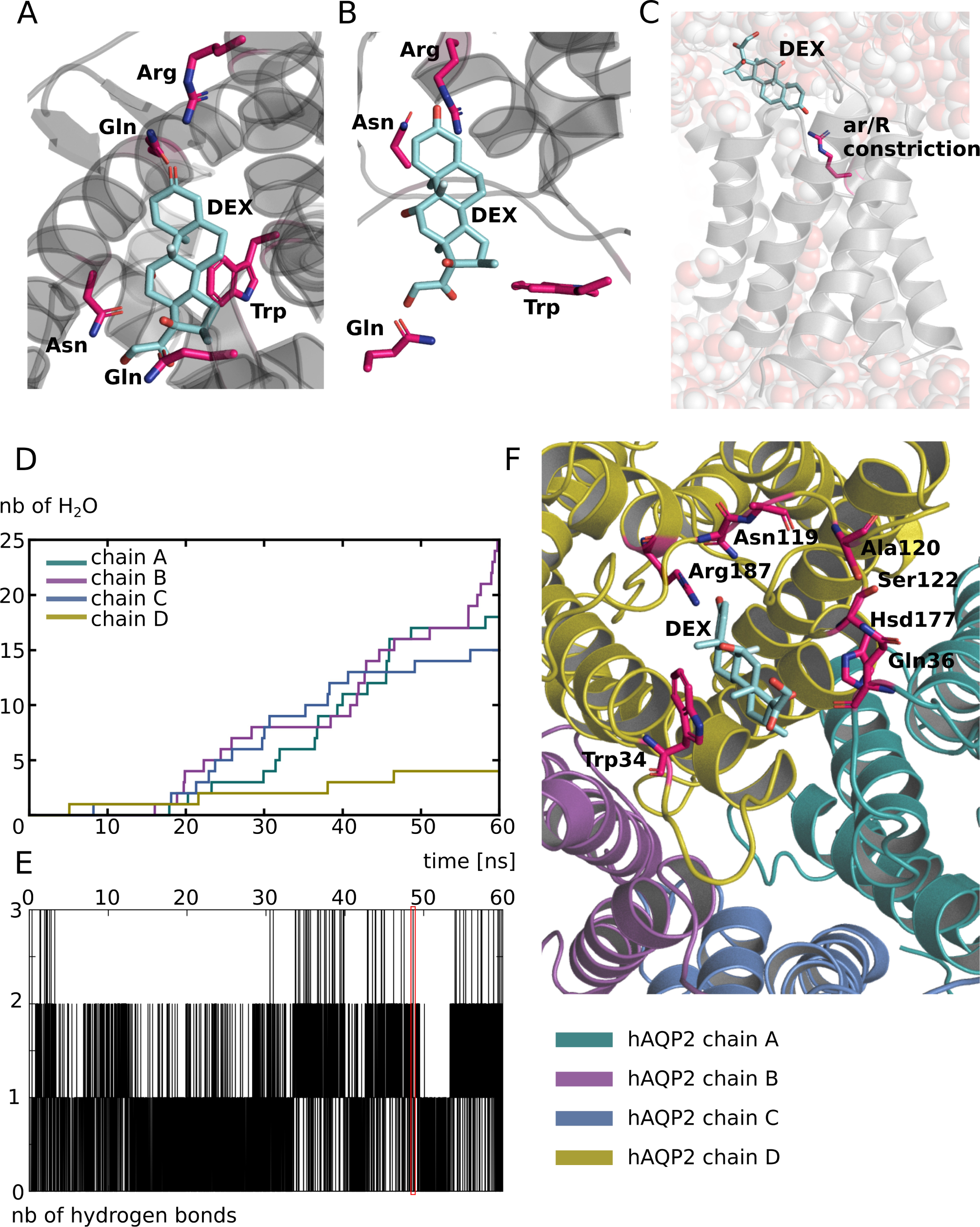
Putative interaction site of dexamethasone with human AQP2. **A**. schematic representation of dexamethasone in its native interaction site. The representation is made form the X-ray structure of the agonist form of mouse glucocorticoid receptor obtained from the protein data bank (pdb : 3mne^44^). **B**. schematic representation of a dexamethasone molecule positioned manually inside the extra-cellular vestibule of human AQP2 (pdb : 4nef) in order to mimic its native interaction with its receptor. Four residues are notably identical between the two proteins dexamethasone interaction sites : an arginine, a tryptophane, an asparagine and a glutamine. These residues were used to guide the manual placement of dexamethasone inside the extra-cellular compartment of AQP2 and are indicated in bold. **C**. Schematic representation of one protomer of the first experimental setup atomic system at simulation time t = 48.6ns. The dexamethasone interacts with the AQP through an hydrogen bond with the arginine of the ar/R constriction. Water molecules are represented with spheres. An interruption of the water molecules continuum is visible. **D**. Cumulative number of water molecules crossing the conducting pore of the channel for each subunit along the 60 ns of simulation. A permeation event is considered when the molecule crosses from end to end a 30 Å long cylinder centered on the center of the channel which corresponds to the whole membrane cross section. **E**. Number of hydrogen bonds between the dexamethasone molecule positioned in the extra-cellular vestibule of chain D and the protein as a function of time. The red box indicates the trajectory frame used as a start point for the other experimental setups and corresponds to simulation time t = 48.6ns. **F**. Schematic representation of a dexamethasone molecule in interaction with AQP2 extracted from the simulation (time = 48.6ns). All of the protein residues involved in the formation of an hydrogen bond along the 60ns of simulation are colored in pink. All representations were made with pymol software^51^.

Then, we simulated the system for 60 ns and observed that the interaction was stable over the whole course of the trajectory, at least for one of the subunits. This is illustrated by the number of hydrogen bonds established between the protein and the dexamethasone molecule positioned in the extra-cellular vestibule of this subunit (chain D) (figure 2.E). All of the residues interacting with the dexamethasone through hydrogen bonding over the 60 ns of simulation are represented on figure 2. F. To get a first impression of the impact of such an interaction on the activity of the channel, we counted the number of water molecules crossing the AQP over the whole transmembrane section (30 Å). The results are displayed on figure 2. D. and clearly indicate a nonfunctional conducting pore for the protomer interacting with dexamethasone over the whole trajectory (chain D). We have the hypothesis that because this interaction includes hydrogen bonding with the arginine of the ar/R constriction (figure 2. F), it will have a significant impact on both criteria impacting the selectivity of the channel (i.e. size and hydrophobicity) and hence on the permeability of the channel. On top of that, the steric hindrance of the dexamethasone itself inside the pore can probably play a significant role as well. This hydrogen bond between the dexamethasone and the ar/R constriction arginine can be visualized on figure 2. C. The interruption of the water molecules continuum is also observed. To further estimate the impact of this putative interaction on the water transport ability of AQP2, a new tetramer is built. We took a conformation of this chain D (which is the protomer presenting the most stable interaction) when the ar/R constriction arginine is involved in an hydrogen bond with the dexamethasone, when the number of hydrogen bonds is the highest (3 simultaneous bonds) and corresponding to the end of the trajectory (at time = 48.6 ns; in order to start from a well relaxed conformation). We then duplicated the protomer in order to reconstitute a tetrameric assembly that we used to further characterize this interaction and its impact on water fluxes through AQP2.

### II. Impact on the transport ability of the channel

#### II.1. Dexamethasone has a significant impact on AQP2 water permeability

To estimate the impact of dexamethasone interaction with AQP2 on water transport, we first compared the whole tetramers with and without dexamethasone together over a 250 ns simulation time course (figure 3). The osmotic permeability coefficients (pf) obtained from the simulations (mean pf of 2.13 .10^−14^ cm^3^.s^-1^ derived from the 250 ns of simulation without dexamethasone) are very close to experimental values (3.3 .10^−14^ cm^3^.s^-1^) issued from heterologous expression systems (constructs expressed in *Xenopus* oocytes)^14^. Moreover, the difference appears very significant between the two conditions (figure 3.A). In a previous work^34^, we highlighted a bias that could be associated to the collective diffusion model used to estimate pf from molecular dynamics simulations^38^. Hence, to strengthen the characterization of dexamethasone impact on water transport, we computed two other permeability estimators as well (figure 3.A). The number of water molecules crossing a 5 Å long section of the channel containing the ar/R constriction constitute a more straightforward approach^34^ and discriminated even more significantly the two conditions (figure 3.A). To be more stringent we also used the number of water molecules crossing the whole conducting pore spanning the entire transmembrane domain over a 30 Å long section and found the same very significant difference between the two conditions (figure 3.A). These first results point toward a significant impact of dexamethasone interaction with AQP2 extra-cellular vestibule on its water transport activity. However, from the cumulative number of water molecules crossing the pore over the 250 ns trajectory, it clearly appears that dexamethasone did not impact the four monomers equally (figure 3.B).

**Figure 3.**
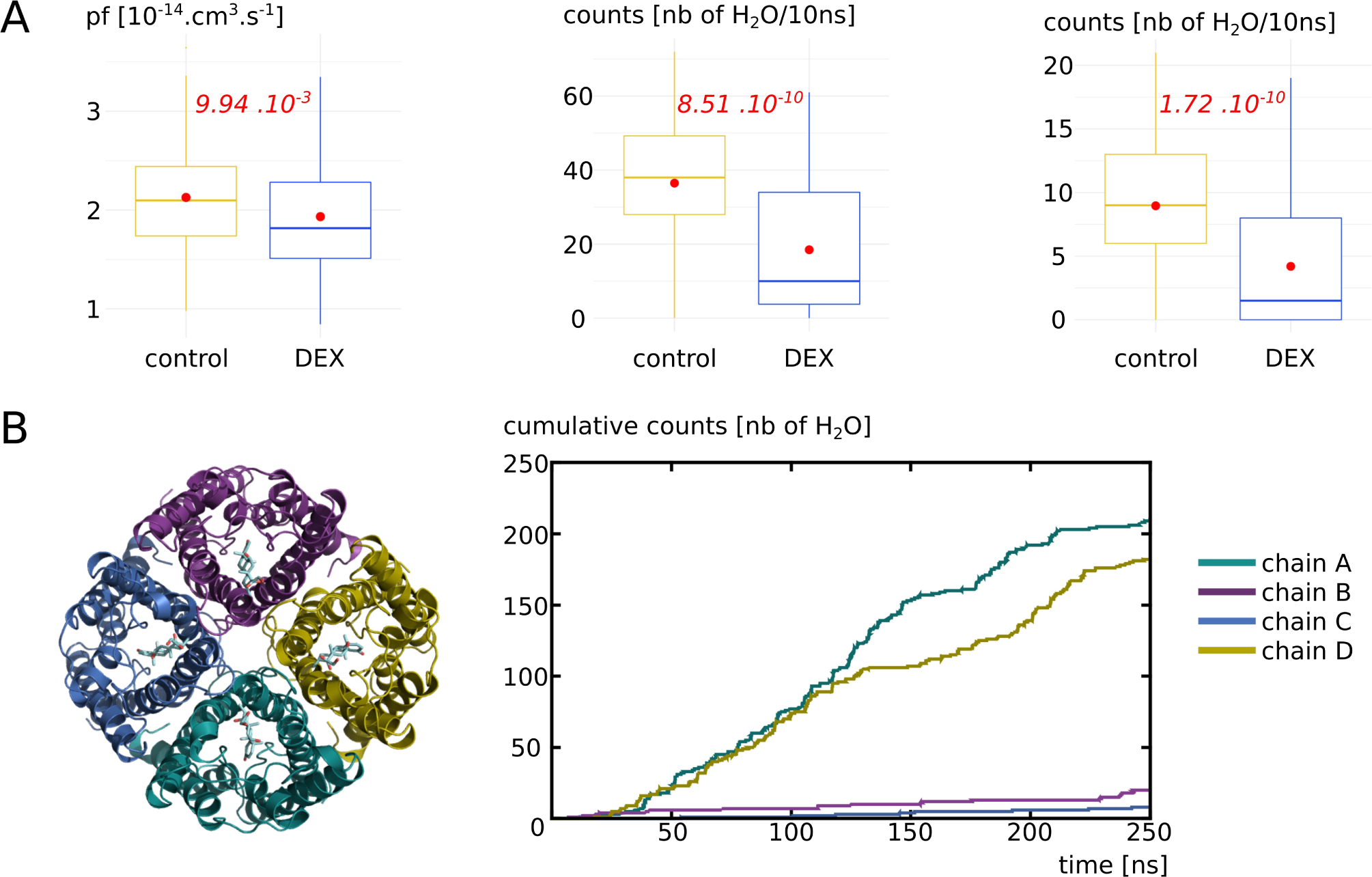
Impact of dexamethasone interaction with AQP2 on the water permeability. **A**. Water permeability is compared between “control” condition (without dexamethasone) and “DEX” condition (with dexamethasone) through different approaches. From left to right : pf derived from the collective diffusion model; number of water molecules crossing a 5 Å section containing the ar/R constriction; number of water molecules crossing the whole conducting pore spanning the entire transmembrane region of 30 Å. Student T test is used to compare pf values and Mann-Whitney test to compare counts. P. values are indicated in red. **B**. On the left, schematic representation of the tetramer of AQP2 with the four dexamethasone molecules positioned in the extra-cellular vestibules at the beginning of the simulation in condition “DEX”. On the right, cumulative number of water molecules crossing the whole transmembrane region of 30 Å for each protomer of the condition with dexamethasone (“DEX”) along the whole trajectory.

#### II.2. Dexamethasone impacts AQP2 water permeability through its interaction with the arginine of the ar/R constriction

To better understand this heterogeneity of response phenotype between the four monomers, we focused our study on the way dexamethasone impacts water permeability in the “DEX” condition (figure 4). We first hypothesized that an high number of hydrogen bonds between the dexamethasone and AQP2 would stabilize the interaction and hence correlate with smaller water permeability. However, we observed the contrary with an high number of hydrogen bonds between the dexamethasone and AQP2 correlating with high water permeability values (figure 4.A). We also hypothesized that an interaction with the arginine of the ar/R constriction would impact the permeability of the channel as it would modify both its size and hydrophobicity. Indeed, we observed a very strong correlation between the smallest distance between this arginine of the ar/R constriction and the dexamethasone and the permeability of the channel (figure 4. B) : the smallest the distance, the smallest the permeability. On top of that, the comparison of pore diameters between the two functional protomers and the two nonfunctional protomers (chain A and D versus chain B and C) yielded a significant difference at the narrowest part of the pore, i.e. at the ar/R constriction region (figure 4. C). Finally, to establish whether the interaction between the arginine of the ar/R constriction and the dexamethasone through hydrogen bonds formation is correlated to the functional state of the pore or not, a Chi2 test is performed. It strongly points toward a statistically significant association between the two variables with a p. value of 4.95 .10^−11^ (figure 4. D). A pore is considered functional based on the precedent set of data (figure 3, see methods section for more information). This association is represented graphically on figure 4. E and F.

**Figure 4.**
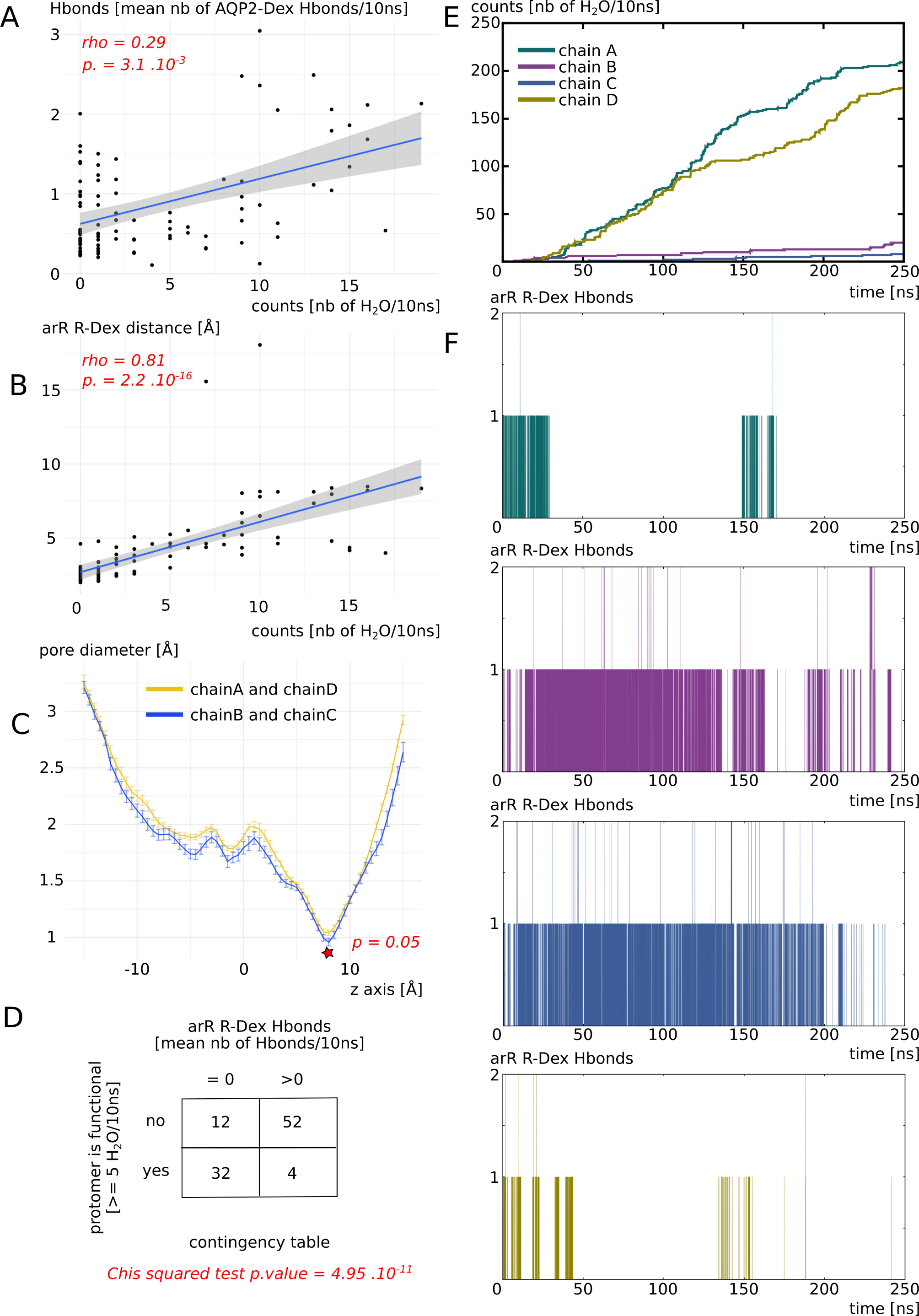
Dexamethasone impacts AQP2 water transport through its interaction with ar/R constriction arginine. All the analysis displayed on this figure were made from the “DEX” condition of figure 3. **A**. Number of water molecules crossing the whole transmembrane section (30 Å long) as a function of the number of Hbonds between each dexamethasone molecule and AQP2. Spearman’s rank rho coefficient is used to estimate the correlation between the two variables. **B**. Number of water molecules crossing the whole transmembrane section as a function of the minimal distance between each dexamethasone molecule and the arginine of the ar/R constriction. Spearman’s rank rho coefficient is used to estimate the correlation between the two variables. **C**. Mean pore diameter for chain A and D (for which dexamethasone leaved the interaction site) and chain B and C (for which dexamethasone stayed stabilized in the interaction site). The standard error is represented. The star indicates the narrowest part of the channel, the ar/R constriction, where a significant difference is found between the two conditions (Mann-whitney test). **D**. Contingency table of the functional state of the protomer (defined as functional when allowing 5 or more water molecules to cross the whole transmembrane section in 10ns) in regards with the presence or absence of Hbonds between dexamethasone and the arginine of the ar/R constriction. **E**. Cumulative number of water molecules crossing the whole transmembrane section as a function of simulation time. **F**. Number of Hbonds between the arginine of the ar/R constriction and dexamethasone for each protomer as a function of simulation time.

Finally, we can conclude that dexamethasone significantly inhibits water transport through AQP2 through its interaction with the arginine of the ar/R constriction (figure 4. D, E and F). This interaction corresponds to a conformational state where the dexamethasone is stabilized close to the ar/R constriction region (figure 4. B), inside the extra-cellular vestibule through a small number of hydrogen bonds with AQP2 (figure 4. A). This interaction directly modifies the size of the pore (figure 4. C). Because the direct interaction (figure 4. D and F) changes the position of the arginine side chain inside the pore and the radiation of the positive charge of the guanidinium, the hydrophobicity of the pore is impacted as well. Hence, Dexamethasone has a significant impact on water permeability of AQP2 through its interaction with the arginine of the ar/R constriction which in turns changes both crucial criteria behind pore water permeability (i.e. size and hydrophobicity^39^).

#### II.3. Detailed impact of dexamethasone interaction with AQP2 on the water permeability as an illustration of pf adjustment with the Dk constant

In a previous work we highlighted a bias that could arise from pf calculation with the collective diffusion method^34^. Due to thermal agitation of water molecules inside the pore, an overestimation of water permeability could be derived from closed channels. To tackle this issue, we proposed a correction constant, the Dk constant. It allows for an higher integration of free energy barriers into pf. Here, we confront this approach to a new set of data. Figure 5 displays the free energy profiles and water permeabilities of each four protomer without (“control” condition) and with (“DEX” condition) dexamethasone. First of all, by simply counting the water molecules crossing the whole transmembrane section, we can confirm the previously observed pattern : chain A and D did not stabilize dexamethasone in the extra-cellular vestibule (see previous section) and are in an open functional state in both conditions while chain B and C through their interaction with dexamethasone (see previous section) are maintained in a closed nonfunctional state in the “DEX” condition.

**Figure 5.**
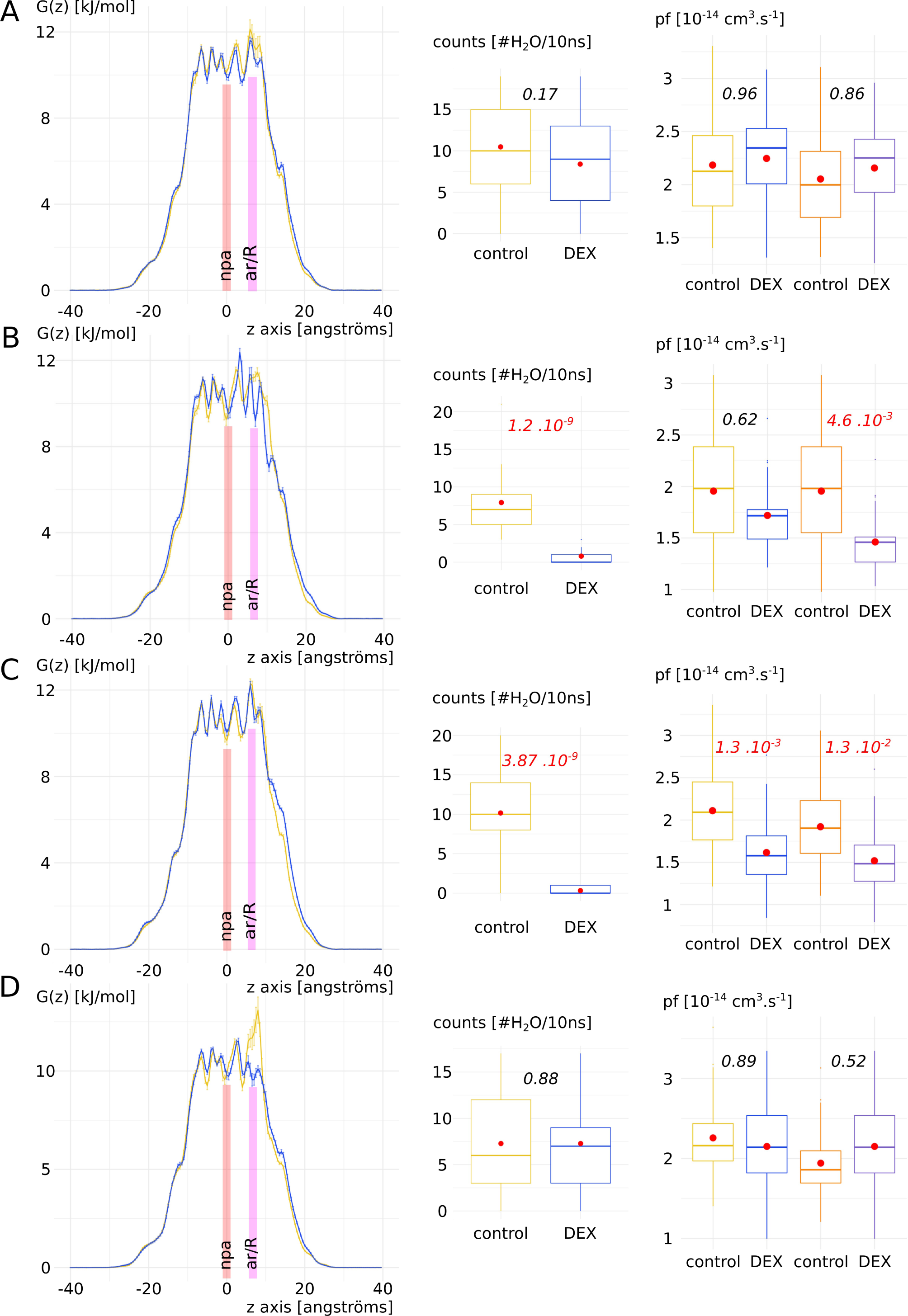
Detailed impact of dexamethasone interaction with AQP2 on the water permeability as an illustration of pf adjustment with the Dk constant. For each chain (**A, B, C** and **D**), from left to right are displayed : free energy profiles, number of water molecules crossing the whole transmembrane section (30 Å long) and the pf without (colors blue and yellow) and with (colors orange and purple) correction with the Dk constant. Control condition without dexamethasone is represented in yellow and “DEX” condition with dexamethasone is represented in blue. For permeabilities comparison between the two conditions, Tukey post hoc test after one-way analysis of variance is used for chains A, C and D and Bonferroni post hoc correction after Wilcoxon test is used for chain B. P. values are indicated in italic.

This pattern is also comforted by the free energy profiles with the apparition of free energy barriers before the ar/R constriction region for chain B and C. An high free energy barrier located at the ar/R constriction is also observed in the control condition of chain D. This could be explained by the propensity of the arginine of the constriction to switch between up and down conformational states^34,35^. This feature has been observed in several AQPs and seems conserved among the family members^34^. AQP2 especially was recently highlighted as particularly concerned by this phenomenon through molecular dynamics study^40^. However, when pf is used to compare the two conditions, significant difference exists only for chain C. We applied the Dk constant correction on pf (see supplementary material and figure S1 for details) and restored the response phenotypes observed through water counts : a significant difference between “control” and “DEX” conditions can be observed for chain B and C. These results add evidence in favor of the Dk correction constant that we introduced in our previous work^34^.

### III. Biological relevance of the dexamethasone – AQP2 interaction

#### III.1. Impact of cation nature on the permeability of AQP2

To test whether the interaction between dexamethasone and AQP2 was likely to happen spontaneously *in vivo*, we built another more realistic atomic system (see methods). This new system takes into account the asymmetric partitioning of ions sodium and potassium between the plasma membrane necessary for the establishment of membrane potential. Hence we placed 150 mM of NaCl in the extra-cellular compartment and 150 mM of KCl in the intra-cellular compartment (see methods, figure 1).

We first estimated the impact of cation nature on the dexamethasone and the AQP. The results are presented on figure 6. Radial distribution functions of sodium, potassium and chloride ions with the dexamethasone molecules as reference show a very quick convergence toward the equilibrium value of 1 which corresponds to a perfect gas state distribution (figure 6.A). To be sure that the dexamethasone molecules were not in interaction with the protein or the membrane and avoid a potential bias, we used the 5 ns of additional equilibration where the dexamethasone molecules are in solution. In contrast, a clear impact of the nature of the cation appears when the reference chosen corresponds to the exposed carboxylates of extra-cellular or intra-cellular surface of AQP2 (figure 6. B). An over-accumulation of cations within a 5 Å range of the protein carboxylates is specific to the sodium but not to the potassium ions (figure 6.B). To explain this phenomenon, we hypothesize that the smallest Van der Waals radius of sodium and its higher electronegativity makes it “stickier” toward negatively charged carboxyl groups.

**Figure 6.**
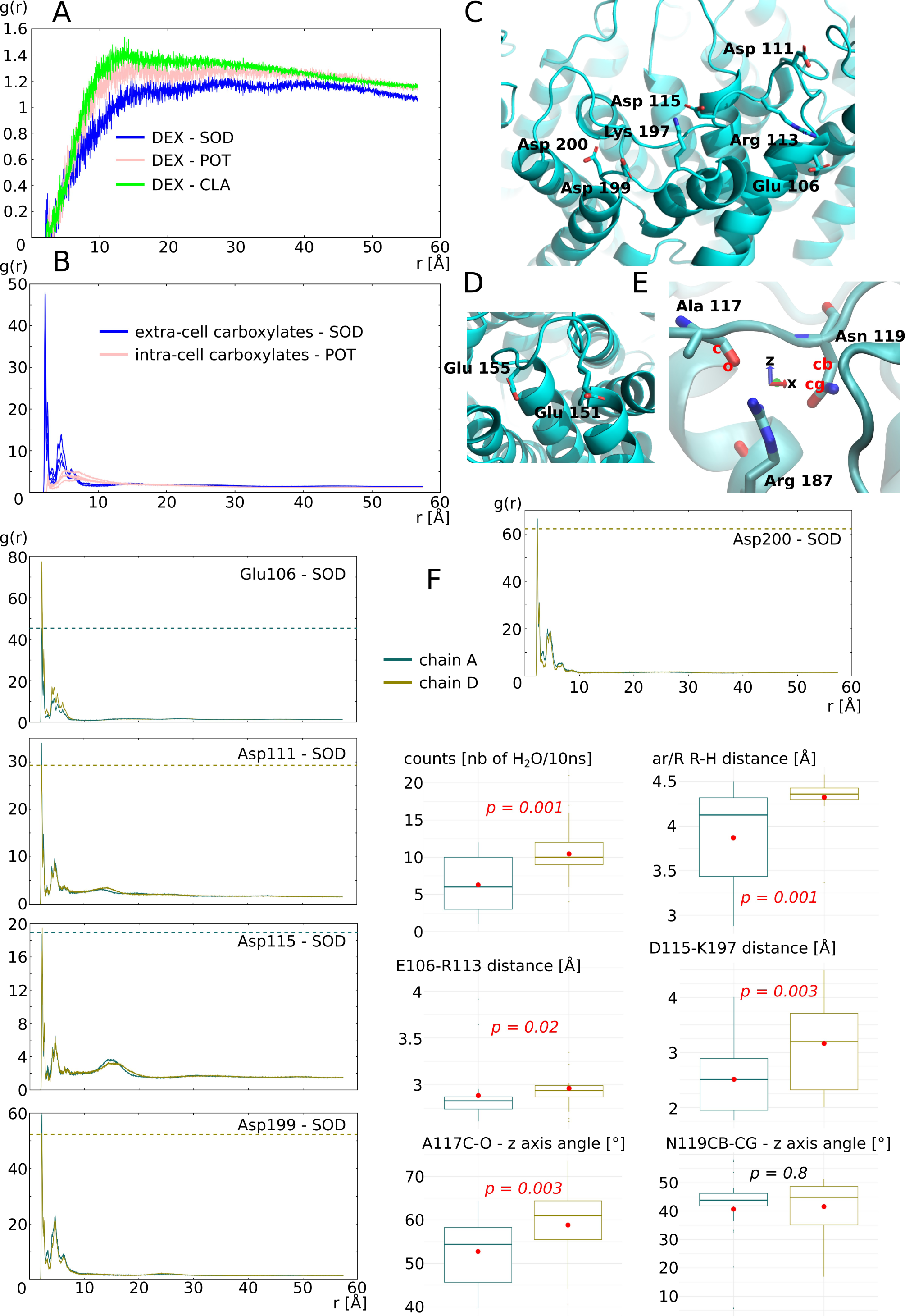
Impact of ion nature on the permeability of the monomers. **A**. Radial distribution function of ions potassium, sodium and chloride with dexamethasone molecules as a reference calculated from the 5ns of equilibration phase. **B**. Radial distribution function of cations with the extra-cellular surface carboxylates or the intra-cellular surface carboxylates as reference. **C**. Schematic representation of the extra-cellular surface carboxylates plus Cloop arginine113 and Eloop lysine 197 involved in salt bridges with these carboxylates. **D**. Schematic representation of intra-cellular carboxylates. **E**. Shematic representation of the arginine of the ar/R constriction with alanine 117 and asparagine 119 of the Cloop with which it forms hydrogen bonds. The angle formed between Ala 117 atoms C, O and the z axis and the angle formed between Asn 119 atoms CB, CG and the z axis are used for further characterization of Cloop torsion. **F**. Characterization of cation nature impact on permeability of the two monomers not interacting with dexamethasone : chain A (in green) and chain D (in yellow). Analysis displayed include : radial distribution function of sodium with each of the extra-cellular surface carboxylate as a reference ; number of water molecules crossing the whole transmembrane section of the channel ; minimal distance between the arginine and the histidine of the ar/R constriction ; minimum distance between Glu 106 and Arg 113 and between Asp 115 and Lys 197 ; angle formed between Ala 117 atoms C, O and the z axis and the angle formed between Asn 119 atoms CB, CG and the z axis. All analysis are performed over the whole 250ns of trajectory. For statistical analysis, chain A and chain D are compared together with non parametric Mann-Whitney test for averaged values over 10ns sub-trajectories i.e. 25 repetitions by chain. P. values are indicated in italic.

Among the four chains of the AQP2 tetramer, two interacted with the dexametasone (chain B and C) and two other did not (chain A and D). We started by characterizing the monomers not interacting with dexamethasone (figure 6. F) : Chain A water permeability is significantly smaller than chain D and this difference correlates with the size of the ar/R constriction. In a previous work, we demonstrated that the electrostatic environment of the ar/R constriction, at the extra-cellular surface of the AQP could impact water permeability by orientating the position of the arginine side chain inside the pore^34^. Hence we hypothesized that the fixation of sodium ions on the carboxylates of the extra-cellular surface could induce this conformational change of arginine side chain reflected by the change in size of the constriction. Therefore, we looked at the radial distribution functions (rdf) of sodium with each extra-cellular carboxylate as a reference separately. We observed an higher over-accumulation of sodium around 3 out of 5 carboxylate (aspartate 111, aspartate 199 and aspartate 200) for chain A compared to chain D. We observed the opposite pattern for glutamate 106 and an equal over accumulation between the two chains for aspartate 115. These observations comfort our hypothesis as an higher number of negative charges located at the periphery of the monomer are attenuated in chain A when compared with chain D and correlates with the difference in water permeability. However, we also observed the complementarity of charged residues positioned on the extra-cellular loops C and E (figures 1.A and 6.C). The Cloop is a mobile loop which dives into the extra-cellular vestibule of AQPs (figure 1.A) and interacts through hydrogen bonds with the arginine of the ar/R constriction implicating alanine 117 and asparagine 119 (figure 6. E). We formulated a complementary hypothesis stipulating that changes of Cloop position mediated by salt bridges formation with Eloop residues could modulate ar/R arginine side chain position through the hydrogen bond network formed between Cloop and the arginine. Based on the residue composition of AQP2, two salt bridges can be formed to stabilize Cloop position : between glutamate 106 and arginine 113 and between aspartate 115 and lysine 197 (figure 6. C). There is an higher over accumulation of sodium at the vicinity of glutamate 106 of chain D which indeed correlates with a significantly higher E106-R113 distance. Aspartate 115 interacts equally with sodium between chain A and chain D. However, it is not the case of aspartate 199 which competes for the interaction with lysine 197 as well. Hence, chain A aspartate 199, by interacting more with sodium than its counterpart in chain D would compete less for the salt bridge formation with lysine 197. This in turn would be in favor of salt bridge formation between lysine 197 and aspartate 115 in chain A reflected by the smallest distance between these two residues. Even though the Cloop stays within a distance allowing for salt bridge formation in both chains (<5 Å), these significant differences still reflect a probable conformational change. Finally, to estimate the impact of such changes on the position and orientation of the Cloop atoms interacting with the arginine of the ar/R constriction, the angle formed between alanine 117 C-O and asparagine 119 CB-CG with the z axis are computed (figure 6. E and F). A significant difference is observed for alanine C-O orientation. This indicates a torsion of Cloop that could have led to the destabilization of hydrogen bond between alanine 117 and arginine 187. Based on all these preliminary results, we can conclude that the sodium ions, through their interaction with AQP2 extra-cellular surface carboxylates, change the charges repartition which in turn impacts the water permeability of the channel through the modulation of arginine 187 side chain position inside the pore. The data presented here are not complete enough to estimate correctly whether the electrostatic surface potential modification, the conformational changes of loopC or both criterion can explain this significant impact of sodium interaction on water permeability of the monomer.

#### III.2. Interaction of AQP2 with dexamethasone without docking

To evaluate better the likelihood of dexamethasone-AQP2 interaction, in this third more realistic setup, the four dexamethasone molecules are placed in solution in both cellular compartments. The dexamethasone interacted very quickly with the AQP2 extra-cellular surface 11 ns after the beginning of the simulation (see movie) and converged toward the predicted interaction site in about 40 ns characterized by the formation of hydrogen bond with the arginine of the ar/R constriction (see previous sections, figure 7. C). We previously showed that the formation of such hydrogen bond was determinant in significantly impacting water permeability (see previous section). Here we observed this phenomenon on two monomers : chain B and chain C (figure 7. C). Two different conformations correspond to this state and differ between the two chains : One implicates the formation of the hydrogen bond with the arginine and the carbonyl oxygen of the dexamethasone (figure 7. G). The other corresponds to the dexamethasone orientated in an opposite manner, presenting its hydroxyl groups toward the guanidinium group of the arginine (figure 7. G). The first conformation was observed inside the extra-cellular vestibule of chain C and inhibits water flux better than the second one, found inside the vestibule of chain B (figure 7. D). It also correlates with a more negative binding free energy (figure 7. F).

**Figure 7.**
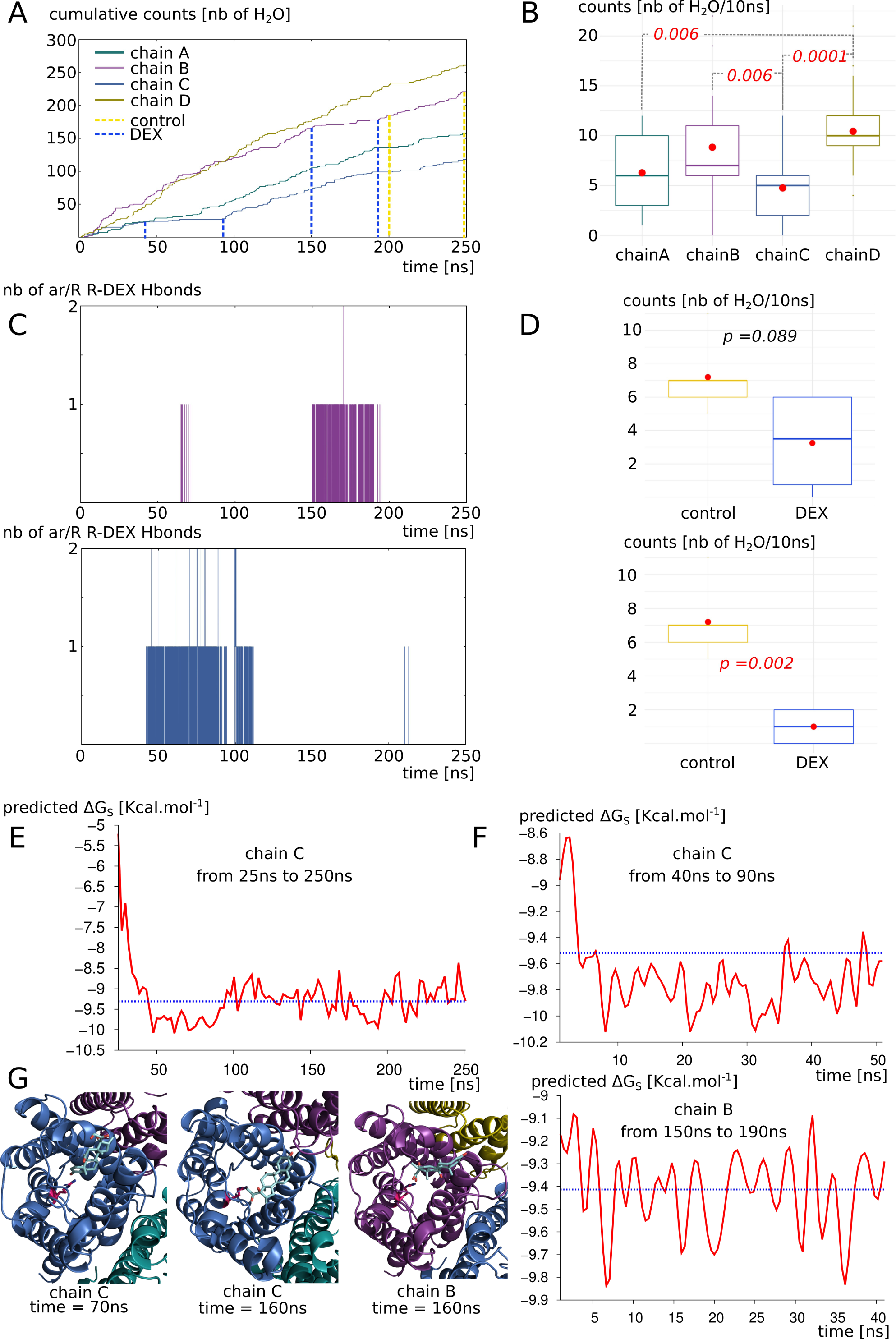
AQP2 interaction with dexamethasone in a biological context mimicking simulation setup. **A**. Cumulative number of water molecules crossing the whole 30 Å long transmembrane section of each monomer for the complete 250ns of trajectory. Dashed lines indicate the sub-parts of trajectory used for statistical analysis of part D and free energy calculation of part F. **B**. Statistical analysis of the water counts (number of water molecules crossing the 30 Å long transmembrane channel per 10ns sub-section of the trajectory). Non parametric Wilcoxon test with post hoc Bonferroni correction is used to compare the four chains. Significant differences are indicated by dashed lines with the corresponding p. value in red italic. **C**. Number of hydrogen bonds formed between the dexamethasone molecules and the arginine of the ar/R constriction of chain B and chain C. **D**. Impact of the fixation of dexamethasone on the protomer permeability (chain B on top and chain C on the bottom). The “control” and “DEX” conditions are issued from the same trajectory but at different times based on the presence of absence of interaction with the dexamethasone (indicated by dashed lines on part A). They correspond to 40 or 50 nanoseconds of simulation. T test is performed to compare conditions. p. values are indicated in italic. **E**. Binding free energy of the dexamethasone 228 with AQP2 with which it mainly interacts with chain C extra-cellular vestibule from time 25ns to the end of the trajectory at time 250ns. Mean free energy is indicated by blue dashed line. **F**. Binding free energy for chain B and chain C sub-sections (40ns and 50ns sections respectively) corresponding to the parts of the trajectory where hydrogen bond is established between dexamethasone and the arginine of the ar/R constriction. Mean free energy is indicated by blue dashed line. **G**. Schematic representation of extra-cellular vestibule of chain B or chain C in interaction with dexamethasone. The arginine of the ar/R constriction is colored in pink.

Even though these conformations impair the water flux the most significantly (see previous section and figure 7. A, C and D), other ones also stabilize the dexamethasone molecule inside the extra-cellular vestibule. Indeed, the dexamethasone 228 interacts with AQP2 inside chain C extra-cellular vestibule from time = 25ns to the end of the trajectory (figure 7. E). During this 225 ns uninterrupted interaction, the dexamethasone switches between conformations (figure 7. G). Even though it did not stabilize the whole time in direct interaction with the arginine of the ar/R constriction, it overall impaired significantly chain C water permeability (figure 7. A and B). The binding free energy corresponding to this interaction with chain C ranges from -5.21 kcal.mol^-1^ to - 10.12 kcal.mol^-1^ with an average of -9.22 kcal.mol^-1^. It corresponds to dissociation constant values (K_D_) ranging from 212004.7 nM to 73.15 nM with an average of 317.7 nM. This last set of data clearly demonstrates the spontaneous nature of the interaction and confirms its significant impact on the water permeability of AQP2.

#### III.3. Proposed molecular mechanism for specific interaction between AQP2 extra-cellular surface and dexamethasone

During the 250 ns simulated, the four dexamethasone of the extra-cellular compartment spontaneously established stable interactions with the AQP2 surface (see movie). In contrast, in the intra-cellular compartment, none of them interacted more than 2 ns with the protein surface. One of them only was quickly stabilized inside the lipidic bilayer and interacted with the hydrophobic transmembrane region of AQP2 for approximately 10 ns (see supplementary figure S2 and movie). We formulated the hypothesis that dipole moment orientation of the molecules coupled with charges attractions could explain this difference. On the upper part of figure 8 are displayed the correlations between the dipole moments of AQP2 tetramer and the dipole moments of the dexamethasone molecule which interacted with chain C extra-cellular vestibule during 225 ns (figure 7. E). And on the bottom part of the figure are displayed the correlations between the dipole moments of AQP2 tetramer and the dipole moment of one of the three intra-cellular dexamethasone not stabilized inside the lipidic bilayer.

**Figure 8.**
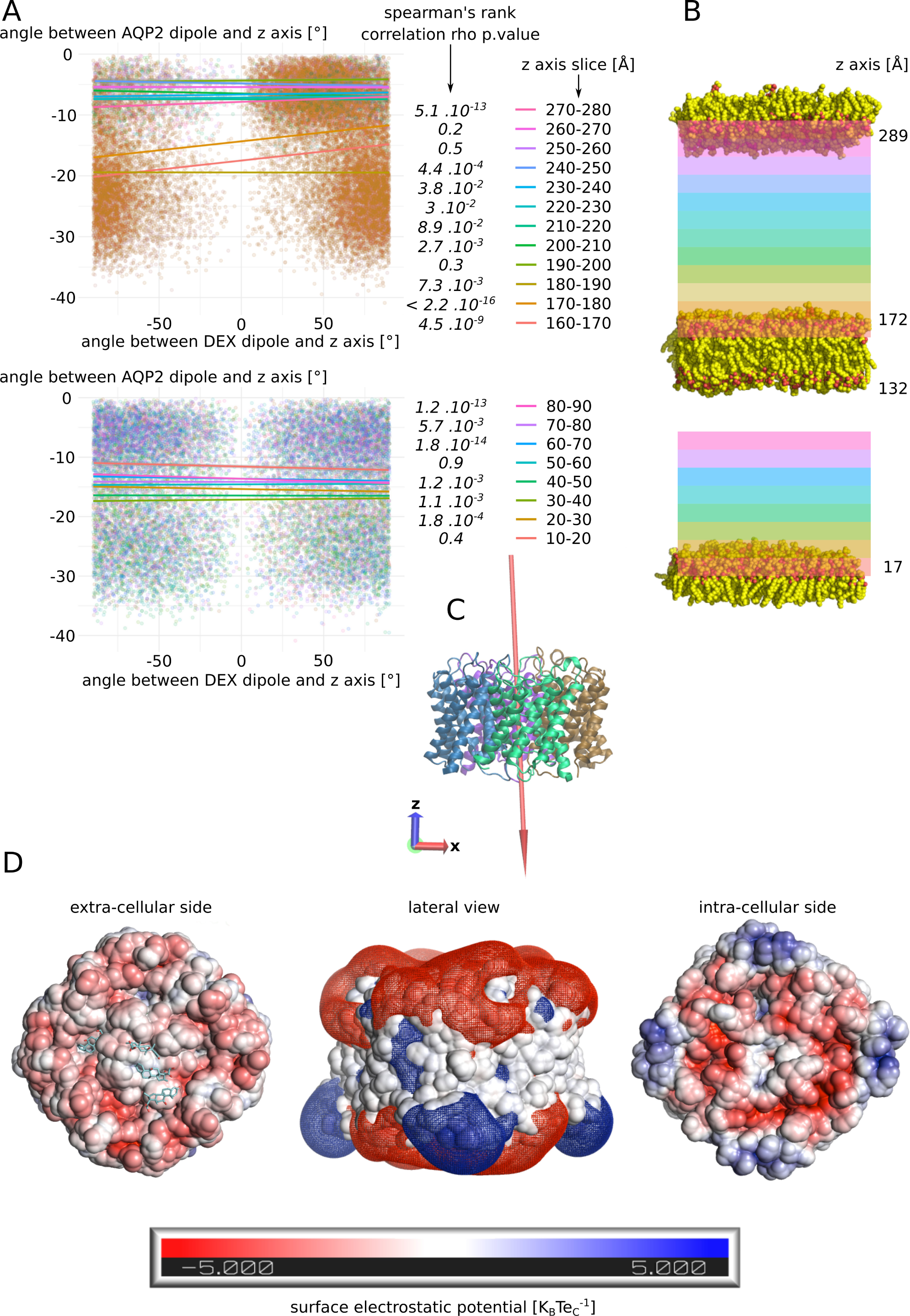
Putative mechanism for spontaneous interaction between AQP2 extra-cellular surface and dexamethasone. **A**. Estimation of the correlation between the dipole moment of the whole tetramer of AQP2 and the dipole moment of the dexamethasone molecule : The angle their dipole moment formed with the z axis of the simulation box is calculated. Then the correlation between angle values for AQP2 tetramer and the dexamethasone is estimated through Spearman’s rank correlation rho. The test is performed according to the position of the dexamethasone molecule in the box. On the upper part, the dexamethasone molecule which interacts with the extra-cellular vestibule of chain C (dex 228) is followed. On the bottom part, a dexamethasone molecule never interacting with the protein (dex 232) is followed. For this later molecule, data above 90 Å along the z axis were missing so the test could not be performed. P. values associated with the correlation tests are indicated in italic. **B**. Schematic representation of the atomic system. The lipids delimiting the two compartments are represented. The z coordinate of their N or O atoms averaged over the whole 250ns trajectory is indicated next to the polar heads. The z axis slices used for correlation tests are schematized with colors corresponding to part A. **C**. Schematic representation of the AQP2 tetramer extracted from the trajectory at time = 250ns. The dipole moment of the tetramer is schematized by a red arrow (scaling = 0.1). The arrow points from the negative pole toward the positive pole. Z and x axis are represented by red and blue arrows respectively. **D**. Surface electrostatic potential of the extracellular and the intracellular faces of AQP2 at time = 50ns. The dexamethasone molecules interacting with the protein are represented in cyan. In the middle, mesh surface of the electrostatic potential is represented (−0.5 K_B_Te_C_^-1^ in red and +0.5 K_B_Te_C_^-1^ in blue).

In order to do so, the dipole moments orientation is compressed inside a single variable chosen as the angle formed between the dipole moment and the z-axis (see methods). The correlation between these angle values computed for AQP2 and the dexamethasone molecule is calculated according to the position of the dexamethasone molecule inside the simulated box. For the extra-cellular compartment, starting from the tetramer surface we can observe a very significant correlation between the dipole moment for the first 20 Å, followed by higher p.values eventually getting to the point of non significant correlations in the upper part of the compartment (slices 250-260 and 260-270) before finally going back to a significant association in the last slice (270-280). These observations could be explained by the influence of the dipole moment of the tetramer (1198.98 debye) upon the smaller dipole (2.93 ± 1.08 debye) moment of the dexamethasone correlated with the charges interactions of the molecule with the protein and the membrane. Indeed, in the first 10 Å, the dexamethasone is interacting directly inside the vestibule of chain C and is hence stabilized in a preferential orientation. In the next slice however, the very low p. value could be the result of a synchronization effect of the small dipole moment of the dexamethasone with the three orders of magnitude higher dipole moment of the tetramer. Then, going further from the protein and higher in the simulated system, this effect would be attenuated by the distance. And finally, the last slice small p. value could be the result of a stabilizing interaction of the dexamethasone with the polar head of the POPC. If the dipole moment of AQP2 orients the dexamethasone molecule, its negative pole would mostly points toward the upper part of the simulated system while its positive pole would be on the contrary mainly orientated toward the bottom part of the system (figure 8. C). This, correlated with the surface charges of the POPC and AQP2, could explain the opposite behavior of the dexamethasone between the two cellular compartments : In the extra-cellular compartment this phenomenon would orient the negative pole of the dexamethasone toward the POPC polar heads displaying their positively charged choline groups and its positive pole toward the negatively charged extra-cellular surface of AQP2 (figure 8. D). In the intra-cellular compartment, the exact opposite situation could be expected as the intra-cellular surface of AQP2 is still mostly negatively charged. This charge repulsion could explain the fact that the correlation is not significant close to the membrane and that there is not enough data to test the correlation near AQP2 surface (figure 8. A and B and movie). Finally, the neutral center of the tetramer probably helps to accommodate the hydrophobic side of the dexamethasone molecules before they reach out one of the monomers vestibule (figure 8. D).

## Discussion

In the present study we have demonstrated the relevance of the AQP2 – dexamethasone interaction. The mean binding free energy corresponds to a K_D_ of 317.7 nM, falling in the range of specific interactions^52^. In the last experimental setup which mimicked a cellular context the best, the dexamethasone placed in solution spontaneously interacted with AQP2, in the short time lapse of 11 ns (figure 7). Moreover, two dexamethasone molecules converged toward the predicted interaction site without external forces to be applied on the system. One of them remained docked inside the extra-cellular vestibule for 225 ns out of the 250 ns simulated (figure 7). This spontaneous interaction was specific to the extra-cellular compartment while in the intra-cellular compartment, no interaction was observed (see movie). To explain this asymmetric behaviour of the dexamethasone relative to its position in the system, we formulated an hypothesis implicating the influence of AQP2 dipole moment on the orientation of dexamethasone coupled with charges attraction or repulsion with the protein and the lipids surface (figure 8). However, more investigations are needed to conclude in favor of this hypothesis. The new interaction site was initially investigated on the basis of the similar features shared with the mouse glucocorticoids receptor X-ray structure (PDB code: 3mne)^44^. The interaction site is located on the extra-cellular surface of AQP2, inside the vestibule of each monomer (figure 2). The tetrameric assembly hence displays four interaction sites for dexamethasone. To estimate the impact of this interaction on the channel water permeability, three different methods were used (pf, number of water molecules crossing the ar/R constriction and number of water molecules crossing the whole transmembrane section: figure 3) in combination with three different atomic systems (initial docking of dexamethasone on the crystallographic structure, reconstructed tetrameric assembly and the same tetramer contextualized inside a compartmentalized system) which all converged toward the same conclusion: there is a significant reduction in water permeability of AQP2 triggered by its interaction with dexamethasone. The dexamethasone reduces water fluxes the most when establishing hydrogen bonds with the arginine of the ar/R constriction (figure 4). This conformation corresponds to the smallest binding free energy (−10.12 kcal.mol^-1^) and the smallest K_D_ (73.15 nM). Even though the interaction between dexamethasone and the arginine of the ar/R constriction is associated with dexamethasone inserted the deepest inside the AQP vestibule, it occurred spontaneously in about 40 ns (figure 7). When the dexamethasone is initially placed in this position (figure 4), we observed the interaction with the arginine maintained stable for 200 ns in two monomers. For the two others, the interaction was observed several times, pointing toward a recurrent phenomenon rather than a rare event. Because the ar/R constriction corresponds to the smallest part of the channel and to an interacting site with water molecules^39,49^, the interaction between the arginine 187 of the ar/R constriction and the dexamethasone impacts significantly both the size and the hydrophobicity of the channel (figure 4). Even though we observed a significant reduction of the pore diameter at the ar/R constriction by dexamethasone interaction, the difference in size is really small (∼0.1 Å, figure 4). Moreover, in the last experimental setup, no significant ar/R constriction size reduction was observed in correlation with dexamethasone fixation (supplementary figure 2). However, the hydrogen bond formed between arginine 187 side chain and the dexamethasone attenuates the positive charge of the guanidinium group, usually radiating inside the pore. This change in electrostatic profile can be estimated through the differences in pore diameter (figure 4) and free energy (figure 5) profiles in other regions than the ar/R constriction. From the free energy profiles, a correction (Dk constant) was applied to pf values, allowing significant differences to emerge between “control” condition and “DEX” condition (figure 5). While the Dk constant was originally designed to integrate better the ar/R constriction free energy barrier into pf^34^, in the present study, the highest free energy barrier used for integration corresponded to the position between NPA motifs and ar/R constriction (supplementary figure 1 and figure 5). Taken together, these results suggest that the main contribution of dexamethasone – AQP2 interaction on water permeability reduction is of electrostatic nature and that the energetic constriction does not correlate with the steric constriction of the channel. We also observed another evidence in favor of the regulatory role of charges distribution in proteins through the impact of sodium on AQP2 water permeability (figure 6). By quenching carboxylates of the extra-cellular surface located on the periphery of the tetramer, sodium ions induced a net conformational change of the arginine of the ar/R constriction conformation characteristic of a closed channel (figure 6). This conformation of the arginine designated as “down state” was already observed in the crystal structures and molecular dynamics studies and linked to the electrostatic environment changes of the protein^34,35,40,53,54^. This result comforts our previous results indicating these extra-cellular carboxylates as playing a significant role in defining the electrostatic environment of the extra-cellular vestibules constituting a relevant regulatory mechanism of AQPs^34^. Adding more weight to this observation, the “down state” of ar/R arginine and its impact on water flux of AQP2 was also described by another team through molecular dynamics study^40^. Even though the authors did not mention the role of ions nature, all their atomic systems were built with 150 mM of NaCl.

Aquaporins have been identified in the inner ear many years ago^55^, in particular in the endolymphatic sac^17,19,21,56^ and are suspected to play a role in the dysregulation of the endolypmh barrier resulting in EH. Already in 1998, AQP2 was suspected to be involved in MD pathogenesis^56^. Different drugs have been thought to alter AQP function, such as anti-diuretic hormone^15,56–58^. AQP2 especially is well known for its role in renal homeostasis under the control of the arginine vasopressin hormone (AVP)^59–61^ and this regulatory mechanism seems conserved in the inner ear as well. Indeed, levels of AQP2 mRNA and water homeostasis of the endolymphatic sac have been shown to be modulated by application of AVP or vasopressin type-2 receptor antagonist (OPC-31260)^15,62–64^ highlighting the regulatory role of AQP2 in the inner ear homeostasis. Other AQPs have been suspected to be involved in the pathogenesis of MD, showing a likely key-role of this type of membrane channels in the pathophysiology of MD^65–67^. Beside the evidence of AQP role in the endolymph homeostasis, corticosteroids (which have been shown effective in the treatment of MD^24–27^) have already been suspected to act through AQP function, in particular by upregulating mRNA of AQP1^68^, or AQP 3 known to be crucial in the reabsorption of water in the inner ear ^69^. Here we bring strong evidence that dexamethasone can also alter AQP2 function in directly impeding water fluxes through this channel, which has never been shown so far, to our best knowledge. This finding thus gives an additional interest to this drug, that may explain part of its efficacy in MD treatment.

First trials of dexamethasone intratympanic administration for MD were conducted in the 1980s^70^ on the premise that the pathophysiology of MD may be immune-mediated^71–73^. Initially prescribed with the aim of reducing potential inflammation within the vestibular endorgans, hypotheses on the mechanisms of action of dexamethasone enriched over time, with modulating actions of epithelial sodium transport^74^, or direct modulation of fluid fluxes^26,28–30^. However, none of these hypotheses have been so far confirmed. A direct action of dexamethasone on the control of water fluxes between inner ear liquid compartments is all the more appealing taken that endolymphatic hydrops has become a hallmark of this pathology^4–6,75,76^. MRI analysis of the inner ear cavities following the so-called “hydrops” protocol (use of contrast enhancers between perilymph and endolymph), now makes it possible to almost systematically correlate increases in the volume of the endolymph, in certain areas of the vestibule, to the symptomology of MD. Therefore pharmacological approaches allowing to prevent or reduce the water accumulation in the endolymph is expected to be beneficial.

The demonstration that a direct molecular interaction between dexamethasone and AQP2 is structurally possible, and that this interaction can significantly alter the fluxes of water through AQP2 constitutes an important step in understanding the possible mechanism of action of dexamethasone in the treatment of Menière disease. It also opens new avenues in the development of novel pharmacological approaches for a better management of the pathology.

## Methods

### Molecular dynamics simulations

All simulations were performed with Gromacs (v.2018.1)^31^ in a CHARMM36m force field^32^. The systems were built with CHARMM-GUI interface^33^. A first minimization step was followed by 6 equilibration steps during which restrains applied on the protein backbone and side chains and on lipids were progressively removed before the production phase performed without restrains. Pressure and temperature were kept constant at 1 bar and 310.15 Kelvin respectively using Berendsen method during equilibration and Parrinello-Rahman and Nose-Hoover methods during production. Lennard-Jones interactions threshold was set at 12 Angströms (Å) and the long-range electrostatic interactions were calculated through particle mesh Ewald method.

Three experimental setups were carried out :

- First, in order to find an interaction site, four molecules of dexamethasone were manually placed in the extra-cellular vestibules of the tetramer (one dexamethasone per monomer) of human AQP2 (PDB code: 4nef, figure 1. A). The dexamethasone molecules were extracted from the structure of a mouse glucocorticoids receptor (PDB code: 3mne) and positioned in the AQP vestibules in order to mimic the native interactions described by this structure. Then the tetramer was inserted into POPC bilayer, solvated with TIP3 water and 150 mM of KCl. The system was then simulated during a time course of 60 nanoseconds (ns). The results corresponding to this first experimental setup are displayed on figure 2.
- Then, we rebuilt a tetrameric assembly of AQP2 by duplicating the subunit of the previous system which interacted with the dexamethasone over the 60 ns of trajectory. The chosen conformation of this subunit corresponds to time = 48.6 ns. This starting conformation was chosen based on two criteria : the high number of hydrogen bonds between the dexamethasone and the protein (3 hydrogen bonds, figure 2.E.) and the presence of the arginine of the ar/R constriction within these hydrogen bonds contributors (figure 2.F). Two tetramers were built this way : one where we kept the dexamethasone from the previous simulation in interaction with the duplicated subunit; and one where we kept the protein only. Two conditions are yielded this way : one with the chosen subunit duplicated into a new tetrameric assembly without dexamethasone called “control” and one with the same tetrameric assembly with four dexamethasone molecules interacting with the four extra-cellular vestibules of the AQP (figure 3.B) called “DEX”. Both tetramers were then inserted into POPC bilayer, solvated with TIP3 water and 150 mM of KCl. Both systems were then simulated during a time course of 250 ns. The results corresponding to this experimental setup are displayed on figures 3, 4 and 5.
- Finally, to estimate the likelihood of the AQP2-dexamethasone interaction further, we built a third system (figure 1.B) : we used the same starting tetrameric assembly as for the second experimental setup condition “control” (i.e. without the dexamethasone fixed inside the extra-cellular vestibules). The tetramer was then inserted into POPC bilayer and solvated with TIP3 water. An additional POPC bilayer without AQP is fused to the initial system to compartmentalize it. Then, in the extra-cellular compartment is added 150 mM of NaCl and in the intra-cellular compartment, 150 mM of KCl. To mimic a membrane potential, an additional NaCl ion pair is placed asymmetrically : the Na^+^ ion is placed in the extra-cellular compartment and the Cl^-^ ion in the intra-cellular compartment^34,35^. As a result, a membrane potential of -30 mV is yielded (figure 1.B). Finally, four molecules of dexamethasone were solubilized into the extracellular compartment and four other into the intra-cellular compartment. The system is then simulated for an additional equilibration time course of 5 ns (without restrains) followed by a production phase of 250 ns. The results corresponding to this experimental setup are displayed on figure 6, 7 and 8.

### Analysis

#### Water permeability

To monitor water molecules displacement along the trajectories, the MDAnalysis library is used^36,37^. From these water coordinates are derived water counts and permeability coefficient (pf). Permeability coefficients were calculated according to the collective coordinate method^38^ (see supplementary data for more details). On figure 5, based on the free energy profiles, a correction (described in our previous work) is applied on pf to integrate better free energy barriers^34^ (see supplementary figure 1 for details).

#### Free energy profiles

Water free energy profiles were extrapolated from the logarithm function of the water counts inside the pore with the z-axis as a reaction coordinate^13,14^. The pore is divided along the reaction coordinate (z axis) in slices of 0.5 Angströms (Å). The average density of water molecules in each slice is then computed over the 250 ns of simulation and the Gibbs free energy G(z) is obtained as follows:

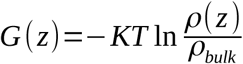

where K, ρ_bulk_ and T represent the Boltzmann constant, the bulk density and the absolute temperature respectively.

#### Binding free energy and dissociation constant

Binding affinity of dexamethasone to AQP2 was evaluated directly from the structure, extracted from the molecular dynamics trajectories every nanosecond, with PRODIGY-LIG program^41^. PRODIGY-LIG evaluates the contacts between ligand and protein and computes a free binding energy from a reliable empirical equation^42^.

Dissociation constant (K_D_) values were obtained from the binding free energies as follow :

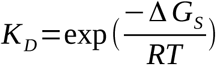

with ΔG_S_, R and T the binding free energy, the perfect gaz constant and the temperature respectively.

#### Other properties

Membrane potential, hydrogen bonds, distances, radial distribution functions (rdf) and dipole moments are computed with gromacs tools (version 2020.6). Pore profiles are computed with HOLE software^43^.

#### Statistical analysis

All statistical analysis are performed using R programming language. Before any statistical test is performed, normality and homoscedasticity of the variables are controlled to choose between parametric or non-parametric tests. When two variables are compared, Student T test or Mann-Whitney test is used. When more than two variables are compared, Tukey post hoc test after one-way analysis of variance or Bonferroni post hoc correction after Wilcoxon test is used.

For experimental setup n°2, the 250 ns trajectories are divided in 10 ns sub-trajectories and the analysis are performed for each monomer hence yielding 100 repetitions per condition for figure 3 and 5 and 25 repetitions per monomer for figure 4. For figure 4.D. contingency table, the functional state of a channel is established as when allowing 5 or more water molecules to cross the whole 30 Å long transmembrane section from one compartment to another in 10 ns. This threshold was defined based on the results of permeability comparison between the condition “control” and “DEX” over the 250 ns of trajectory and displayed on figure 3. The value of 5 water molecules was elected as a functional state threshold because it discriminates both means and medians of the two conditions.

For experimental setup n°3, the 250 ns trajectory is divided in 10 ns sub-trajectories and the monomers are compared between each other yielding 25 repetitions per condition for figures 6 and 7.B. On figure 7. D., 40 ns or 50 ns slices of trajectory only are used for analysis, hence yielding 4 to 5 repetitions per condition. On figure 8, the system is divided in 10 Å long slices according to the z axis. To estimate the correlation between the dipole moment of the tetramer and the dipole moment of the dexamethasone, the angle they form with the z axis is used. The z axis is chosen as reaction coordinate because of the native stable orientation of AQP2 tetramer dipole moment almost parallel to the z axis (figure 8. C). For each slice, the frames in which the center of geometry of the dexamethasone considered (residue number 228 for extra-cellular compartment and residue number 232 for intra-cellular compartment) was contained into the slice were used to extract the dipole moment of the dexamethasone and the corresponding dipole moment of AQP2 tetramer. The Spearman’s rank rho coefficient is then used to estimate the correlation between angle values for each slice.

## Abbreviations

MD: Menière’s disease
EH: endolymphatic hydrops
AQP2: aquaporin 2
Å: Angströms
PDB code: Protein Data Bank code
POPC: 1-Palmitoyl-2-oleoyl-sn-glycero-3-phosphocholine
TIP3: transferable intermolecular potential 3
mM: milliMolar
KCl: potassium chloride
ns: nanoseconds
pf: permeability coefficient

## Supplementary

### pf calculation and correction with the Dk constant

The single-channel permeabilities (*pf*) were computed independently for each protomer based on the collective diffusion model proposed by Zhu et al^1^. Accordingly *pf* was computed from the slope of the mean-square displacement (MSD) of the collective water coordinate. The slope was computed by fitting a line to the MSD up to a time displacement of 100 ps. For each protomer, the analysis was carried out on water molecules within a cylinder of length 8 angströms, centered on the two NPA motifs asparagines Cα center of geometry, that is within the narrowest part of the channel where water molecules are known to form a single file continuum.

In a previous work^2^, we introduced a correction constant to accentuate the effect of the ar/R constriction calculated from the free energy profiles as follows:

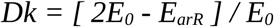

with *Dk* the unit free correction constant ; *E*_*arR*_ the free-energy at the ar/R constriction site and *E*_*0*_ the free energy corresponding to the highest free-energy barrier in the channel section used to calculate *pf. E*_*0*_ must be smaller than *E*_*arR*_ for the correction to be applied. *Dk* integrates the contribution of the ar/R constriction to water diffusion and is comprised between 1 and 0 : when the difference between the two free-energy barriers tends toward 0, *Dk* tends toward 1. On the other hand, the higher the free-energy barrier of the constriction is, the smaller *Dk* is, eventually reaching a limit of the correction when *E*_*arR*_ becomes more than twice as high as *E*_*0*_. In this case, *Dk* becomes negative and is considered as equal to 0. To adjust the *pf*, one has to multiply it by *Dk*:

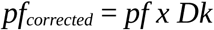

In the present study, we broaden this *Dk* correction to other free energy barrier than the ar/R constriction free energy barrier only. Hence, *E0* still corresponds to the highest free-energy barrier included in the channel section used to calculate pf. And *E*_*arR*_ is replaced by *E*_*HB*_ (*HB* standing for Highest Barrier) and designate the highest free energy barrier outside of the channel section used to calculate *pf*:

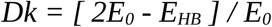

This approach was used to compute corrected pf values of figure 5 and is detailed on supplementary figure 1.

**Supplementary figure 1.**
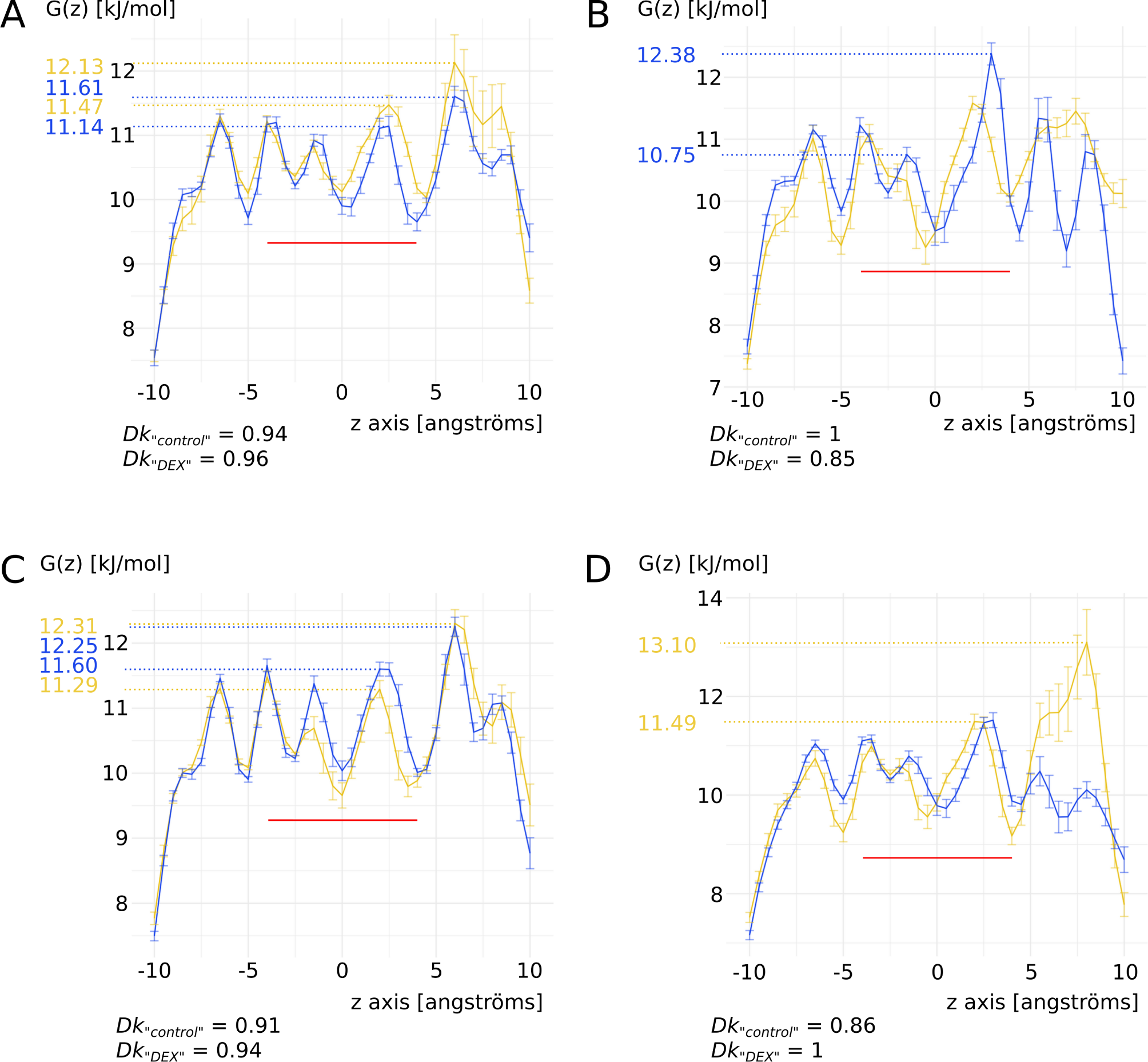
Dk constant for pf correction. For each chain (**A, B, C** and **D**) is displayed free energy profiles of water zoomed on the 20 Å of the water channels. The z axis coordinates are relative to the center of geometry of the Cα of the two asparagines of the NPA motifs (i.e. the center of the conducting pore). The two conditions of the second experimental setup “control” (in yellow) and “DEX” (in blue) are compared. The red line indicates the region used for pf calculation. The free energy values used as E_0_ and E_HB_ for Dk constant calculation are indicated by dashed lines.

### The energetic barrier does not always correlate with the steric barrier

Figure S2 illustrates the absence of correlation between water counts and size of the ar/R constriction between the four chains in the third experimental setup (see methods). The conformational and charge repartition changes induced by sodium interaction have a clear impact on the size of the ar/R constriction through their impact on arginine 187 side chain position inside the lumen of the pore of chain A. However, the modulatory effect of dexamethasone binding on chain B and chain C is more subtle and impacts water flux through modifying the pore electrostatic profile rather than by simple steric occlusion (see results). More experiments need to be performed to shed more light on the molecular mechanisms involved.

**Supplementary figure 2.**
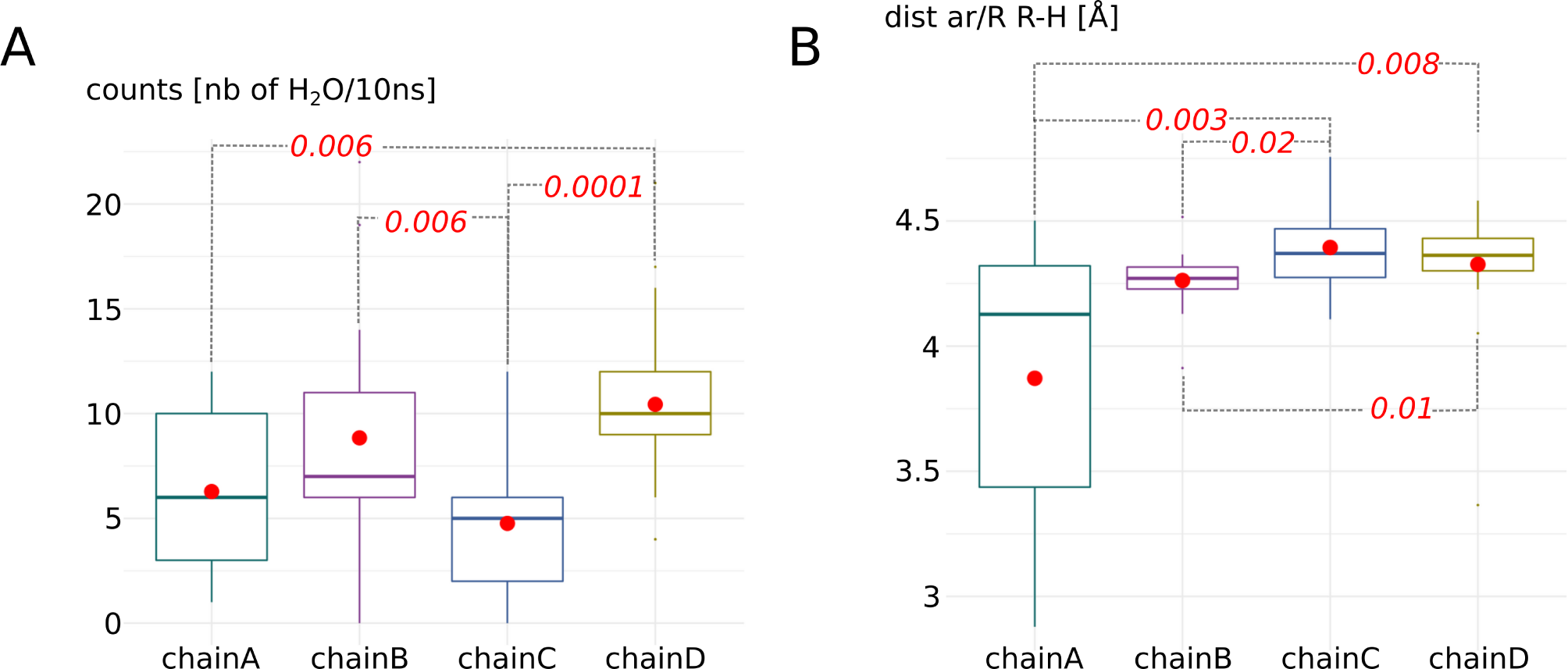
Water channels permeabilities do not correlate with minimal ar/R R-H distances in third experimental setup. **A**. Statistical analysis of the water counts (number of water molecules crossing the 30 Å long transmembrane channel per 10ns sub-section of the trajectory). Non parametric Wilcoxon test with post hoc Bonferroni correction is used to compare the four chains. Significant differences are indicated by dashed lines with the corresponding p. value in red italic. **B**. Statistical analysis of the minimal distance between the arginine and the facing histidine of the ar/R constriction. Non parametric Wilcoxon test with post hoc Bonferroni correction is used to compare the four chains. Significant differences are indicated by dashed lines with the corresponding p. value in red italic.

### Dexamethasone stabilization in POPC bilayer

In the last experimental setup, we observed spontaneous stabilization of dexamethasone inside the POPC bilayer (figure S3). For both compartment, the dexamethasone was stabilized inside the upper membrane (relatively to the box coordinates). This is in favor of our hypothesis stipulating that AQP2 dipole moment orients dexamethasone molecules inside the simulation box. If true, this would result in the dexamethasone molecules presenting their negative pole toward the upper membrane where the positively charged choline groups of POPC would constitute good interacting partners. On the opposite, no stabilization event of dexamethasone into the lower membranes occurred during the 250 ns of simulation. However these preliminary data are not sufficient to conclude.

**Supplementary figure 3.**
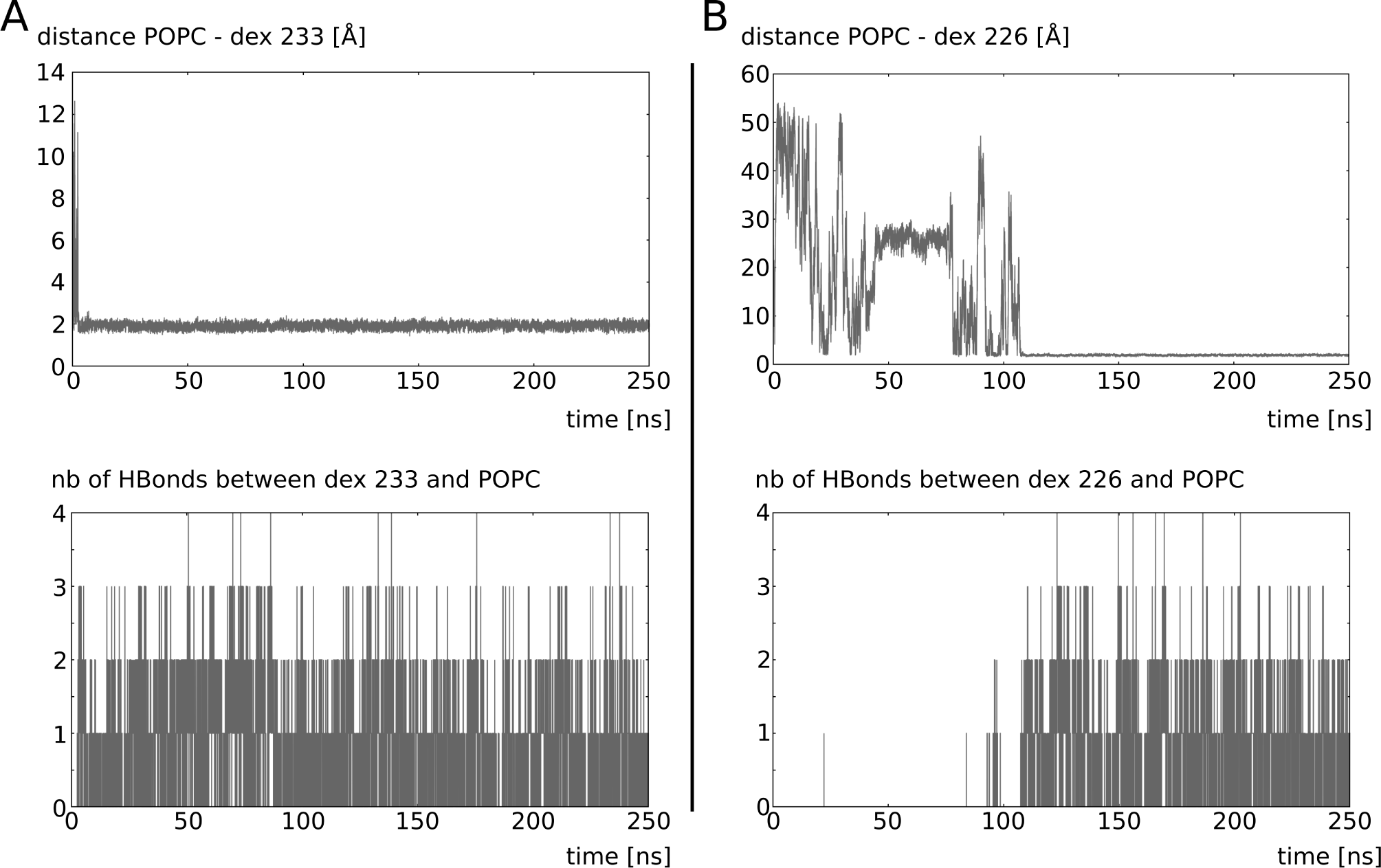
Dexamethasone stabilization in POPC bilayer. **A**. Dexamethasone residue 233 of intra-cellular compartment and **B**. Dexamethasone residue 226 of extra-cellular compartment. For both molecules, the minimal distance between the dexamethasone and the POPC is displayed as a function of time as well as the number of hydrogen bonds established between them as a function of time. The data are extracted from the third experimental setup (see methods).

